# Impact of bead-beating intensity on microbiome recovery in mouse and human stool: *Optimization of DNA extraction*

**DOI:** 10.1101/2020.06.15.151753

**Authors:** Bo Zhang, Matthew Brock, Carlos Arana, Chaitanya Dende, Lora Hooper, Prithvi Raj

## Abstract

DNA extraction methods play an important role in the acquisition of accurate and reproducible 16S sequencing data in microbiome studies. In this study, we assessed the impact of bead-beating intensity during DNA extraction on microbiome recovery in mouse and human stool. We observed a higher DNA yield, better DNA integrity, higher *Shannon’s entropy* and *Simpson’s index* in samples beaten for 4 and 9 minutes as compared to unbeaten samples. 16S sequencing data showed that bead beating has a statistically-significant (p<0.05) impact on the recovery of many clinically relevant microbes that live in the mouse and human gut, including *Bifidobacterium, Sutterella* and *Veillonella.* It was observed that 4 minutes of bead beating promotes recovery of about 70% of OTUs in mouse and human stool, while the remaining 30% requires longer bead beating. In conclusion, our study indicates adjustments in bead beating treatment based on the composition of the specimen and the targeted bacteria.

## Introduction

High throughput sequencing technology is commonly used to characterize microbial composition of biological specimens. This approach can be applied to capture microbial diversity in human and environmental specimens with unprecedented depth (1–4). A number of prior studies provide evidence that methods of sample collection, storage and DNA extraction are critical for accurate profiling of microbiota in environmental (5–7) or human samples (8–10). In particular, it is increasingly apparent that the DNA extraction method is crucial to the accuracy of microbiome analysis (11–13). Given that the microbial composition of a niche is generally diverse with significant variations in cell membrane structures and functions among community members, obtaining a complete and unbiased representation of microbial DNA from all community members is technically challenging.

There is growing evidence that complete lysis of bacterial cell walls is critical for optimum yield of DNA. Lysis protocols include procedures that lead to physical and or enzymatic disruption of the microbial cell wall (5, 14, 15). It has been observed that extended lysis time and mechanical disruption can enhance nucleic acid yield. However, extended lysis time can also reduce molecular complexity by shearing genomic DNA into smaller fragments (16, 17). In general, bacterial cells are lysed to release the nucleic acids and the remaining proteins are discarded. Gram-positive bacteria pose the greatest challenge for complete lysis due to their thick cell walls and complex cell wall composition, consisting of several layers of peptidoglycan (18).

Given that the precise composition of pathogenic clinical specimens is mostly unknown and may vary significantly from sample to sample, an ideal DNA extraction method should accurately recover DNA from a wide variety of bacteria and avoid the bias that can be introduced by incomplete cell wall lysis. Bead-beating is a method of mechanical disruption that is performed prior to standard DNA extraction. In this step, ceramic or glass beads are added to the tube containing microbial samples. This is followed by moderate to high speed shaking, causing collisions between the beads and the samples. Bead-beating has become a common method of bacterial cell lysis in microbial metagenomics studies, and a number of different bead beating protocols have been used to extract microbial DNA from stool samples (19). Here we have assessed the impact of bead-beating time on extraction efficiency of nucleic acids and abundance and composition of bacterial OTUs in mouse and human stool.

## Materials & Methods

### Sample collection

We collected two mouse (C57/Bl6) stool samples, designated WT1 and WT2, and two human stool samples, designated Hum1 and Hum2. The stool samples were collected under sterile conditions and stored in DNA/RNA shield, a nucleic acid stabilizing solution from Zymo Research (R1100). DNA/RNA shield provides an accurate molecular signature of the sample at the time of collection by preserving nucleic acids at ambient temperature and inactivating organisms including infectious agents.

### DNA extraction

We used the ZymoBIOMICS™ DNA Miniprep Kit (D4300) for DNA extraction on both mice and human stools. Figure 1 illustrates the experimental workflow of the study. Each of the mouse and human stool samples was aliquoted into four subsamples for the experiment. About 200 mg of feces was aliquoted into a ZR BashingBead lysis tube (0.1 and 0.5 mm). For lysis, 750 ul of ZymoBIOMICS lysis solution was added to each sample tube. Next, each sample tube was tightly closed and loaded onto the PowerLyzer 24 Homogenizer (110/220 V) from Qiagen for bead beating. WT1 and WT2 and Hum1 and Hum2 were two independent replicates of mouse and human feces, respectively. We selected four different bead beating time points as illustrated in Figure 1: 0 minutes (no bead-beating at all), 1 minute (one cycle of shaking), 4 minutes (2 cycles of 2 minute shaking, with a 30 second pause after each cycle) and 9 minutes (4 cycles of 2 min and 1 cycle of 1 minute, with a 30 second pause after each cycle). Each of these samples were bead-beaten at a speed of 2200 RPM and were maintained at a temperature of 20°C throughout the bead beating process. Following beat-beating and lysis, DNA was purified using the ZymoBIOMICS protocol, and 100 ul was eluted for downstream experiments. The DNA concentration was measured using the Picogreen method (Invitrogen Quant-iT™ Picogreen dsDNA Assay Kit Reference No. P11496 on Perkin Elmer 2030 Multilabel Reader Victor X3) and DNA integrity number (DIN) was determined on 4150 Tapestation from Agilent using Agilent’s gDNA Screen Tape (Reference No. 5067-5365) and Agilent’s gDNA Reagents (Reference No. 5067-5366).

**Figure 1.**
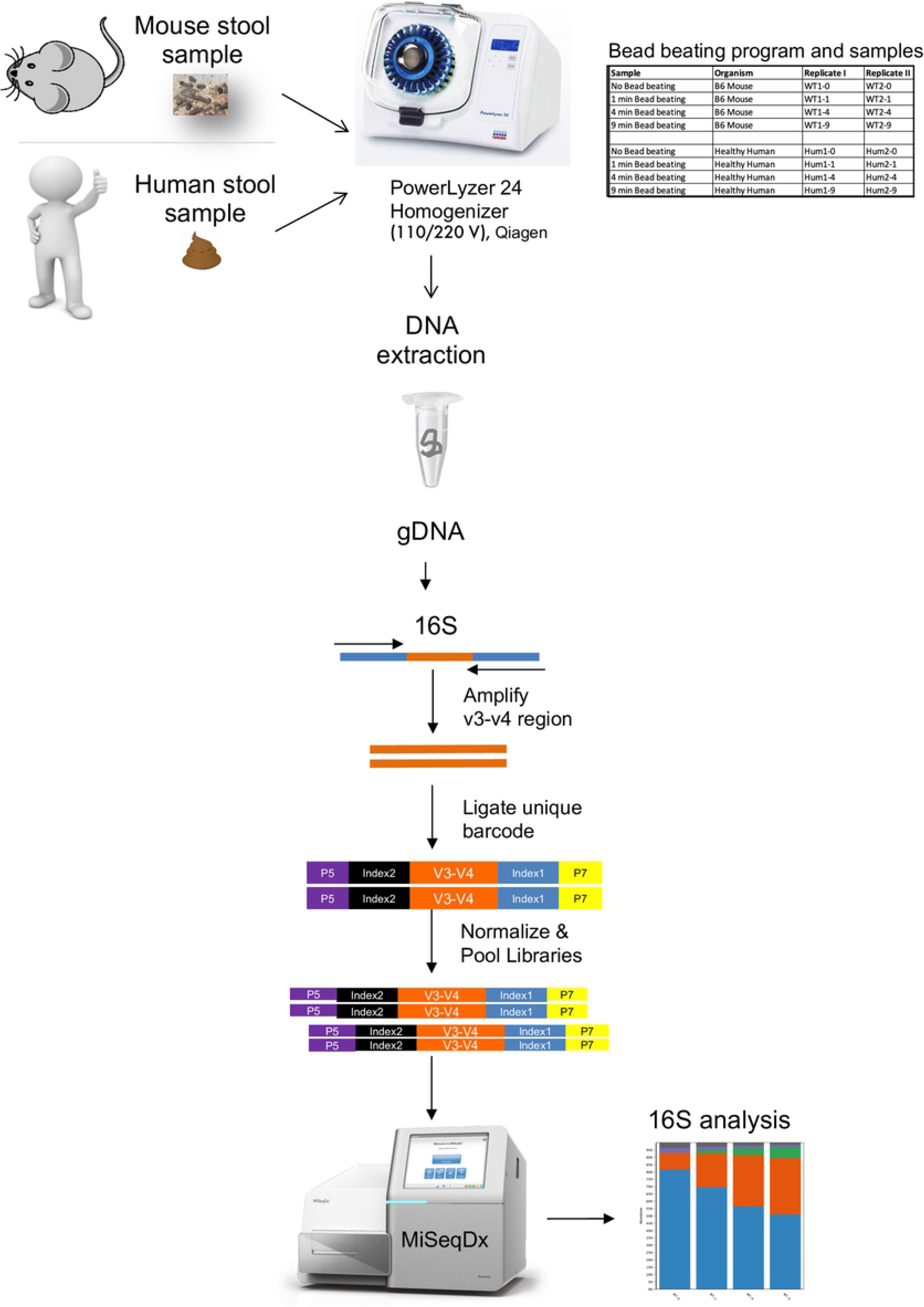
Experimental workflow for 16S sequencing. Illustration of the experimental workflow. Two mouse and two human stool samples were homogenized using a PowerLyzer 24 Homogenizer (110/220V; Qiagen). DNA was extracted using four different bead beating times, followed by 16S rRNA gene sequencing and analysis.

### 16S rRNA gene sequencing

10-50 ng of purified DNA from stool was used to amplify hypervariable region V3-V4 of the bacterial 16S rRNA gene using the Illumina Nextera protocol (Part # 15044223 Rev. B). A single amplicon of about 460 bp was amplified using the 16S Forward Primer **(5’** TCGTCGGCAGCGTCAGATGTGTATAAGAGACAGCCTACGGGNGGCWGCAG) and the 16S Reverse Primer **(5’** GTCTCGTGGGCTCGGAGATGTGTATAAGAGACAGGACTACHVGGGTATCTAATCC) as described in the Illumina protocol. The PCR product was purified using Agencourt AmpureXP beads from Beckman Counter Genomics. We used the Nextera XT Index Kit v2 (Reference no. 15052166) for 16S amplification. Illumina adapter and barcode sequences were ligated to the amplicon in order to attach them to the MiSeqDx flow cell and for multiplexing. Quality and quantity of each sequencing library were assessed using Bioanlyzer and picogreen measurements, respectively. The libraries were then pooled in equal concentrations according to picogreen measurements. Each pool was quantified using KAPA Biosystems Library Quant Kit (illumina) ROX Low qPCR Mix (Reference No. 07960336001) on an Applied Biosystems 7500 Fast Real-Time PCR system. According to the qPCR measurements, 6 pM of pooled libraries was loaded onto a MiSeqDX flow cell and sequenced using MiSeq Reagent Kit v3 600 Cycles PE (Paired end 300 bp). Raw fastq files were demultiplexed based on unique barcodes and assessed for quality.

### 16S data analysis pipeline

Samples with more than 50K QC pass sequencing reads were used for downstream 16S OTU analysis. Taxonomic classification and Operational Taxonomic Units (OTUs) abundance analysis were done using the CLC Bio microbial genomics module (https://www.qiagenbioinformatics.com/plugins/clc-microbial-genomics-module/). Individual sample reads were annotated with the Greengene database and taxonomic features were identified. Alpha and beta diversity analysis was done to understand within- and between-treatment group diversity, respectively. Raw fastq files from this study have been submitted to the Sequence Read Archive with ID PRJNA625828.

## Results

### Assessment of DNAs extracted using different bead beating times

First, we measured the amount of total DNA recovered from each bead-beating treatment. As expected, the bead-beaten samples yielded higher amounts of DNA as compared to unbeaten samples. As shown in Supplementary Fig.1 A-B, the highest yields were observed in samples beaten for 4 or 9 minutes. The DNA integrity number (DIN) was highest in samples treated for 1 and 4 minutes (Supplementary Fig.1C-D). The number of pass filter sequencing reads was highest in mouse stool samples that were beaten for 4 and 9 minutes (Supplementary Fig.1E). However, in human stool samples, the highest pass filter reads were obtained at the 1 and 4-minute time points (Supplementary Fig.1F). We also compared the total number of high-confidence OTUs annotated in all the samples. As shown, the highest OTUs were observed in samples beaten for 4 or 9 minutes (Supplementary Fig. 1G-H). Overall, 4 minutes of beating time was found to give the optimum results for all the assessed parameters.

### *Actinobacteria* requires extensive bead beating for maximal recovery

QC pass sequencing reads were used to define OTUs (operational taxonomic units) at different taxonomic levels such as phylum, class, order, family, genus, and species (Fig. 2A-B, Supplementary Table S1-S4). 16S analysis showed that *Actinobacteria* were significantly (p<0.05) underrepresented in unbeaten samples. Their maximal recovery was observed after 4 and 9-minutes of bead-beating (Fig.2C-D). On the other hand, *Proteobacteria,* which are Gram-negative organisms, were better captured in unbeaten samples or after just 1 minute of bead beating (Fig. 2I&J). Bacteroidetes were least affected by bead-beating time in both mouse and human stool samples (Fig. 2G&H). Results for *Firmicutes* were not consistent between mouse and human samples, as more *Firmicutes* were recovered at 4 and 9 minutes of bead-beating of mouse stool whereas no such trend was observed in the human samples. The aggregated phylum level abundances and comparative statistics between time points in mouse and human data are given in Supplementary Tables S3 and S4, respectively. Differential abundance analysis revealed OTUs that differed significantly between 0, 1, 4 and 9-minutes of bead-beating of mouse and human stool (Supplementary Tables S5 & S6). Supplementary Tables 7A&B list genus level annotations of top OTUs in mouse and human stool.

**Figure 2.**
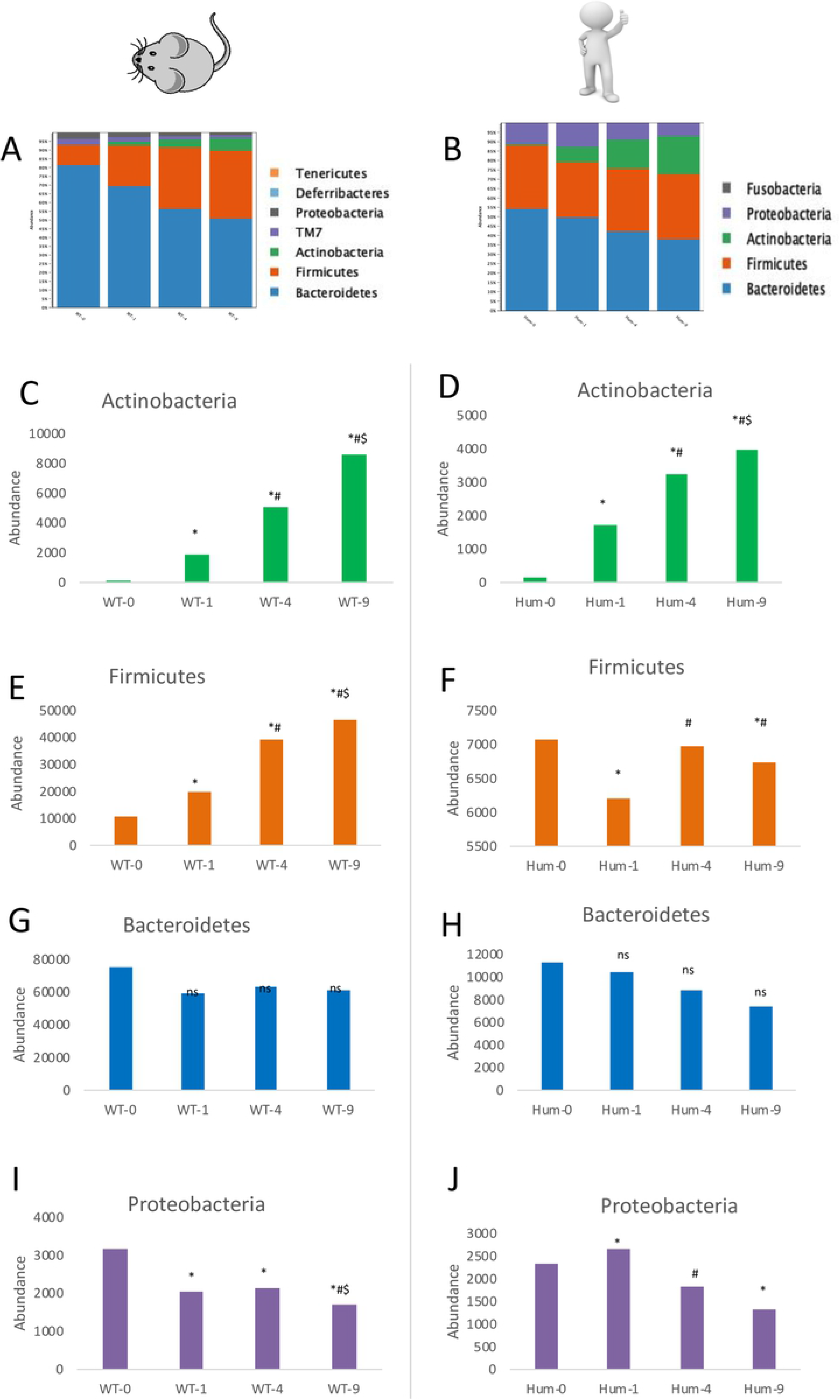
*Actinobacteria* are strongly impacted by bead beating in mouse and human stool. Panels A-B: Color coded bar plots showing the phylum level abundance across different bead beating treatments in mouse and human stool, respectively. Panels C-D show abundance of *Actinobacteria* across bead beating treatments in mouse and human stool, respectively. Panels E-F show abundance of *Firmicutes* in mouse and human stool beaten for different times. Panel G-H shows abundance of *Bacteroidetes* in mouse and human stool. Similarly, in Panels I-J, bar plots show abundance of *Proteobacteria* across four bead beating time points in mouse and human stool. Statistical p-values are denoted with *, # and $ represent comparison with samples that were unbeaten, or beaten for 1 minute and 4 minutes, respectively.

### High bacterial diversity in bead beaten samples

Alpha diversity analysis showed higher phylogenetic richness in bead beaten samples as compared to unbeaten samples (Supplementary Fig.2A, E). Shannon’s entropy and Simpson’s indices are metrices that are commonly used for measurement of bacterial diversity. As shown in Supplementary Fig. 2B &F, higher Shannon entropy was observed after 1, 4 and 9-minutes of bead beating as compared no bead beating. Similarly, Simpson’s indices were higher in bead-beaten samples, further suggesting high bacterial recovery at 4 and 9 minutes of bead beating (Supplementary Fig. 2C&G). As shown in Supplementary Tables S8 & S9, bead beaten samples showed a 1.1-fold increase in phylogenetic diversity, *Simpson’s index* and *Shannon entropy* as compared to unbeaten sample. Beta diversity analysis showed that all bead beaten samples clustered more closely to one another than to unbeaten samples (Supplementary Fig. 2D &H). Overall, it was observed that most of the diversity was captured by beating for 4 minutes and no significant increase in diversity was noticed with further bead beating.

### Bead beating duration strongly impacts recovery of clinically relevant bacteria

Differential abundance analysis on the most abundant OTUs revealed five clusters of bacteria (Fig. 3A, Supplementary Table S10). As shown, cluster 1 (C1) was comprised of *Bifidobacterium* and *Ruminicoccus* in human stool. Maximum recovery of these bacteria was observed after 4 and 9 minutes of beating as compared to no bead beating (C1 in Fig.3A). On the other hand, abundance of *Sutterella*, *Veillonella dispar* and *Veillonella parvula* DNA was highest in samples that were unbeaten or beaten for 1 minute as compared to samples beaten for 4 or 9 minutes (C2 in Fig. 3A). Another cluster of bacteria in human stool was comprised of *Blutia obeum*, *Bifidobacterium longum*, *Coprococcus, Dorea* and *Streptococcus*. These organisms were more highly represented at the 4-minute timepoint and did not show a significant increase in recovery with longer bead beating (i.e., 9 minutes). Cluster 4 (C4) was comprised of *Lactobacillus reuteri*, *Allobaculum* and *Bifidobacterium pseudolongum* in mouse stool. Maximum abundance of these bacteria was observed after 9 minutes of bead beating (Fig. 3A, C4). On the other hand, bacteria of the *Rikenellaceae, Desulfovibrio, Bacteroidales* and *Clostriadales* groups showed maximum abundance in unbeaten samples, as shown in cluster 5 (C5) of Fig. 3A.

**Figure 3.**
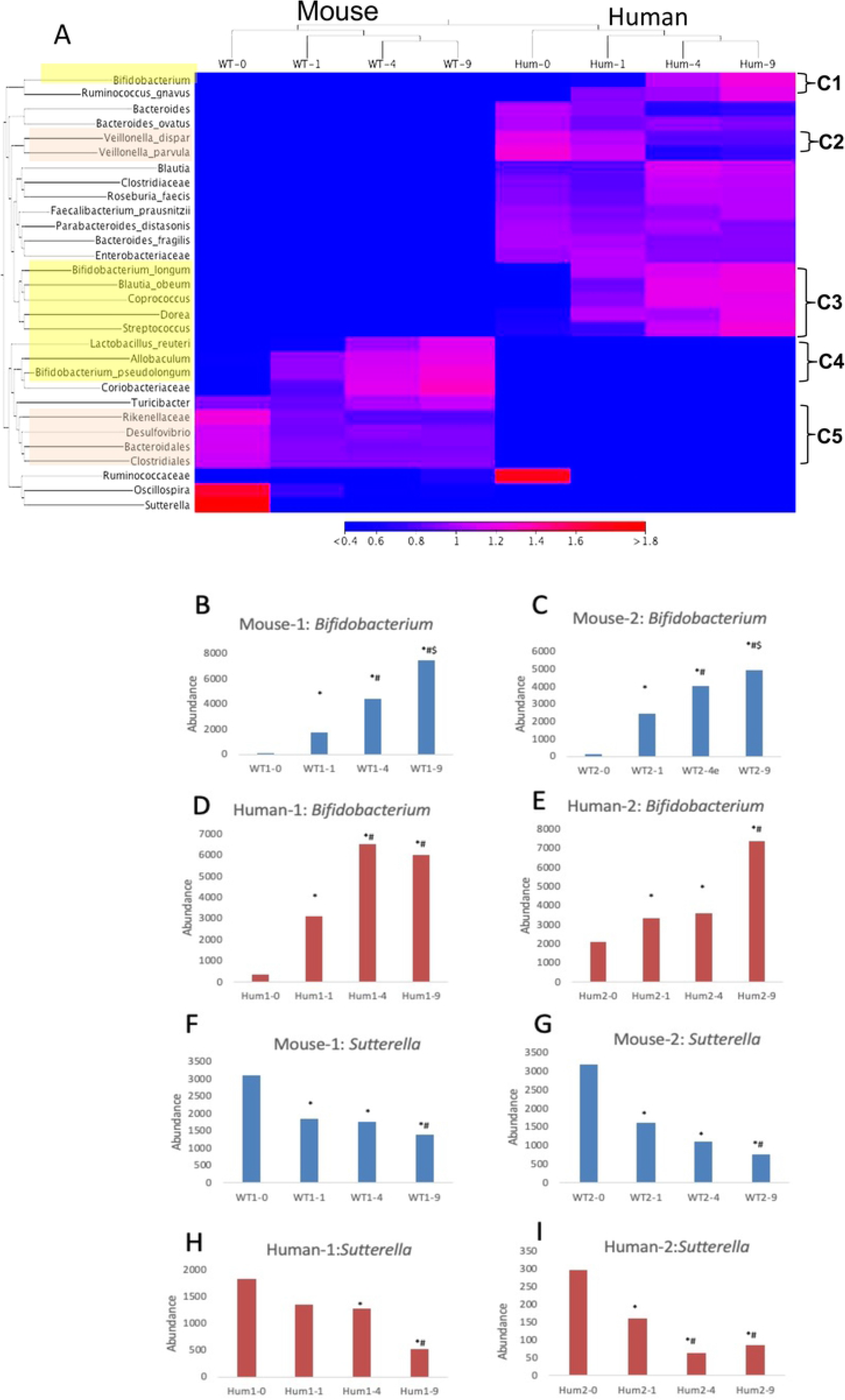
Bacterial clusters defined by bead beating time. Panel A: Results of differential abundance analysis. The heatmap shows the top 30 differentially recovered OTUs in mouse and human stool. Panels B-E show the abundance of *Bifidobacterium* across four beating treatments in mouse and human stool. Similarly, Panels F-I show the abundance of *Sutterella* across four beating treatments in mouse and human stool. Data from replicates of mouse and human sample is presented. Statistical p-values denoted with *, # and $ represent comparison with samples that were unbeaten, or beaten for 1 minute and 4 minutes, respectively.

Interestingly, we found that bead beating intensity has a strong impact on the recovery of clinically-relevant inhabitants of mouse and human gut, including members of the genera *Bifidobacterium, Sutterella* and *Veillonella.* As shown in Fig. 3B-E, replicates of mouse and human stool showed maximum abundance of *Bifidobacterium* in samples beaten for 9 minutes, with 30-100-fold higher recovery in mouse and 2-16-fold higher recovery in human stool upon bead beating. On the other hand, maximum abundance of *Sutterella* was observed in mouse and human stool samples that were unbeaten or beaten for the least amount of time (Fig. 3F-I). We observed a 2-4-fold reduction in *Sutterella* abundance in bead beaten stool, suggesting an adverse effect of beating on recovery of DNA from this bacterial group. These results were consistent across mouse and human stool replicates (Fig. 3F-I, Supplementary data in Table S11-12).

### Optimum bead beating time for maximal recovery of microbiome diversity

We compared various parameters including nucleic acid yield, DNA integrity, sequencing depth and OTU counts across beating times in order to determine the optimum beating intensity for mouse and human stool analysis. We found that optimum data were obtained with 4 and 9 minute bead beating treatment as compared to no bead beating or beating for 1 minute. Comparison of samples beaten for 4 and 9 minutes did not show marked differences. In data from mouse stool, there were only 7 OTUs (out of 24 major OTUs) whose abundance differed significantly (p<0.05) between samples beaten for 4 and 9 minutes. These were *Bifidobacterium*, *Adlercreutzia, Allobaculum, Coriobacteriaceae, Lactobacillus, Turicibacter* and *Ruminicoccus* (Supplementary Table S7A-B). Similarly, *Streptococcus, Suttrella, Dorea, Parabacteroides* and *Bifidobacterium* were 5 of 27 major OTUs in human stool that differed significantly (p<0.05) in samples beaten for 4 versus 9 minutes. These results suggest that up to 70% of microbial signatures can be captured with just 4 minutes of bead beating. However, stool samples rich in bacteria such as *Bifidobacterium*, *Streptococcus* and *Adlercreutzia*, etc. may require more than 4 minutes of beating for maximal DNA recovery. These results suggest that 4-5 minutes of bead beating may be sufficient to capture most of the bacterial diversity in mouse and human stool.

## Discussion

In this study we have systematically assessed the impact of bead beating on microbiome analysis of mouse and human stool. Due to multiple technical and environmental factors, an accurate and reproducible characterization of microbiota composition is a major challenge. Methods of sample storage and collection, DNA extraction, sequencing library preparation and bioinformatics analysis have been shown to contribute variability in 16S results (20–24). Of these, the DNA extraction method is among the most important in that it can introduce bias at the initial step.

Several studies have reported optimization of DNA extraction methods and have developed protocols for extracting microbial DNA from stool samples (8, 9). Large scale microbiome studies such as Human Microbiome Project (HMP), MetaHIT, and the Earth Microbiome Project have reported improved versions of DNA extraction protocols for various types of samples (25–27). The published literature suggests that complete lysis of bacterial cell walls using beads can markedly impact DNA yield as well downstream 16S sequencing results (28, 29). Observed maximal recovery of *Actinobacteria* in samples subjected to bead beating for 9 minutes is consistent with published literature that reports enhanced nucleic acid recovery from Gram-positive organisms with longer disruption of the bacterial cell wall (30). However, there are also other factors such as volume and temperature of elution buffer, type of lysis beads, lysis tubes and columns that were not evaluated in the current study but can also impact overall DNA yield and sequencing data quality.

Our data suggest that bead beating duration strongly impacts the recovery of DNA from several groups of bacteria. For example, optimization of the duration of bead beating enhanced DNA recovery from *Bifidobacteria*, *Sutterella* and *Veillonella*, three clinically-relevant bacterial groups that are important members of the mouse and human gut microbiome (19, 31–35). *Bifidobacterium,* a genus that is significantly underrepresented in the analysis of unbeaten stool, is one of the major colonizers of the human gastrointestinal tract. These microbes have been shown to provide health benefits to their host and are investigated in the context of various human diseases such as colorectal cancer, necrotizing enterocolitis and inflammatory bowel diseases (31).

By contrast, we found that recovery of DNA from certain bacterial groups was reduced by bead beating. For example, DNA from *Sutterella* and *Veillonella* showed reduced prevalence in samples after bead beating, suggesting sensitivity of these microbes to extensive mechanical lysis. These bacteria are also clinically relevant, as altered abundance of *Sutterella* has been associated with many clinical conditions such as autism spectrum disorder, down syndrome and inflammatory bowel disease (32, 33). Similarly, epidemiological studies in young children have associated *Veillonella* with asthma (34), bronchiolitis (36) and autism (35). Since abundance of these microbes could be clinically informative, it is important to be able accurately and precisely determine their abundance in clinical specimens. Our data suggest that studies targeting *Bifidobacteria* should incorporate longer (up to 9 minutes) bead beating protocols in order to ensure maximal recovery of DNA from these bacteria, while those targeting organisms such as *Sutterella* and *Veillonella* should avoid extensive bead beating for maximal recovery and accurate representation. Our data indicate that 4-5 minutes of bead beating may be appropriate to process samples where the composition of microbiomes are unknown.

In summary, our study demonstrates that the duration of bead beating has a strong impact on the recovery of DNA from clinically relevant microbiota in both mouse and human gut. Our data suggest that a minimum of 4 minutes of bead beating (using Qiagen PowerLyzer) can result in recovery of about 70% of gut microbiota DNA signatures. Further, our study identifies particular groups of bacteria in mouse and human stool that can be recovered with up to 4 minutes of bead beating and those that require extensive bead beating for maximal recovery.

## Acknowledgments

This study was supported by the UT Southwestern Microbiome Research Laboratory. Authors gratefully acknowledge donors of deidentified human stool samples for the study.

## Author contributions

B.Z. and M.B. performed experiments; C.A. performed sequencing quality control, C.D. collected mouse stool for study, L.V.H. contributed to manuscript editing and P.R. conceived and designed the experiments and wrote the manuscript.

## Data availability

Raw fastq files from mouse and human experiments have been deposited in NCBI SRA database with accession no. PRJNA625828

